# Mechanisms of scaffold-mediated microcompartment assembly and size-control

**DOI:** 10.1101/2020.10.14.338509

**Authors:** Farzaneh Mohajerani, Evan Sayer, Christopher Neil, Koe Inlow, Michael F. Hagan

**Affiliations:** Martin Fisher School of Physics, Brandeis University, Waltham, Massachusetts, USA; Department of Biochemistry, Brandeis University, Waltham, Massachusetts, USA

## Abstract

This article describes a theoretical and computational study of the dynamical assembly of a protein shell around a complex consisting of many cargo molecules and long flexible scaffold molecules. Our study is motivated by bacterial microcompartments, which are proteinaceous organelles that assemble around a condensed droplet of enzymes and reactants. As in many examples of cytoplasmic liquid-liquid phase separation, condensation of the microcompartment interior cargo is driven by long flexible scaffold proteins that have weak multivalent interactions with the cargo. We describe a minimal model for the thermodynamics and dynamics of assembly of a protein shell around cargo and scaffold molecules, with scaffold-mediated cargo coalescence and encapsulation. Our results predict that the shell size, amount of encapsulated cargo, and assembly pathways depend sensitively on properties of the scaffold, including its length and valency of scaffold-cargo interactions. Moreover, the ability of self-assembling protein shells to change their size to accommodate scaffold molecules of different lengths depends crucially on whether the spontaneous curvature radius of the protein shell is smaller or larger than a characteristic elastic length scale of the shell. Beyond natural microcompartments, these results have important implications for synthetic biology efforts to target new molecules for encapsulation by microcompartments or viral shells. More broadly, the results elucidate how cells exploit coupling between self-assembly and liquid-liquid phase separation to organize their interiors.

---

While the eukaryotic cytoplasm is organized by lipid-encased organelles, it is now clear that cells from all kingdoms of life employ other modes of compartmentalization, such as liquid-liquid phase separation [1–4] and proteinaceous organelles [5–7]. For example, bacterial microcompartments are organelles that form by assembling a protein shell around a dense complex of enzymes and reactants in certain metabolic pathways[8–15]. The best-studied type of microcompartment is the carboxysome, which assembles around rubisco and carbonic anhydrase to enable carbon fixation in cyanobacteria [8, 9, 16, 17].

Other protein-shelled compartments are found in bacteria and archea (e.g. encapsulins [18, 19] and gas vesicles [18, 20]), and eukaryotes (e.g. vault particles [21]). Microcompartment function depends crucially on the amount and composition of encapsulated cargo, and the integrity of the surrounding shell. Thus, understanding the mechanisms that control cargo encapsulation and shell assembly are important to elucidate the normal biological functions of microcompartments, and to reengineer microcompartments as customizable nanoreactors for synthetic biology applications (e.g. [12, 22–35]).

More broadly, there are strong parallels between the condensed cargo within a microcompartment and liquid-liquid phase-separated fluid domains. For example, the rubisco complex within a carboxysomes has a similar structure and function as the pyrenoid [36, 37]. Moreover, experiments and models suggest that microcompartment assembly proceeds by phase separation of the cargo, either prior to or during assembly of the outer shell [38–46]. However, the protein shell of a microcompartment confers distinct advantages over uncoated liquid domains. In addition to functioning as a selectively permeable barrier [47–49], experiments suggest that the shell plays a structural role, influencing the size and morphology during microcompartment assembly [23, 26, 38, 50–52]. Despite intense recent investigations into LLPS, we have yet to understand the role of LLPS in promoting or guiding self-assembly and cargo encapsulation, or conversely the role of assembly in promoting and regulating LLPS. Thus, models that describe coupled LLPS and microcompartment assembly are needed to identify the factors that control microcompartent formation, cargo encapsulation, and function. From a practical perspective, condensation of cargo into fluid domains is a powerful approach to develop self-assembling and self-packaging gene/drug delivery vehicles [53–63]. Coupling such complexes to shell assembly could provide a means to control their size, amount of packaged cargo, and targeting capabilities.

Microcompartment shells are polydisperse (40-600 nm diameter), roughly icosahedral, and formed from pentameric, hexameric, and pseudo-hexameric (trimeric) protein oligomers [8, 9, 17, 51, 64–68]. Recently, Sutter et al. obtained atomic-resolution structures of small (40 nm diameter) empty microcompartment shells composed of pentamer, hexamer, and pseudo-hexamer proteins [51] or smaller shells (25 nm, 21 nm) composed of only pentamer and hexamer proteins [68]. While these structures provide essential insight into microcompartment architectures, the effect of cargo on shell assembly, and the mechanisms that control cargo packaging, remain unclear. In many types of microcompartments, cargo coalescence and cargo-shell interactions are mediated by auxiliary microcompartment proteins called scaffold proteins. Scaffold proteins are typically flexible (e.g. the α-carboxysome scaffold (CsoS2) is an intrinsically disordered protein) and contain multiple interaction sites with cargo molecules [37,41,43]. Thus, scaffolds drive cargo coalescence via weak multivalent interactions, paralleling mechanisms that drive liquid-liquid phase separation in cells.

Previous modeling studies have elucidated the assembly of empty icosahedral shells [69–95], the effect of an interior template such as a nanoparticle or RNA molecule on shell assembly[76, 82–84, 96–118], and how the interplay between template curvature and shell elasticity can control its size [76, 104, 105]. However, the many-molecule scaffold-cargo complex within a microcompartment does not have a specific size or morphology, and thus requires new models. Recent computational works investigated assembly of microcompartment shell subunits [119, 120] or assembly of shells around a single cargo species with direct pairwise attractions driving cargo coalescence and packaging [44–46]. These studies showed that the cargo can influence the size of the assembled shell through a kinetic mechanism[46] or a combination of kinetic and thermodynamic effects [44, 45]. However, there are no models for microcompartment assembly that account for the crucial role of scaffold molecules.

Therefore, in this work we develop a computational model for microcompartment assembly that explicitly accounts for flexible scaffold molecules and scaffold-mediated cargo coalescence and packaging. We are motivated by recent experiments that investigated how changing the length of the scaffold molecule affects the size of *α*–carboxysome shells [43, 121], but the model is sufficiently minimal and general to also provide insights into self-assembling delivery vehicles (e.g. [53–62]). We also develop an equilibrium theory, which extends models for spherical micelles assembled from star or diblock copolymers[122–126], to describe variable packaging of the interior block, and an exterior shell with a preferred curvature radius (which may differ from the preferred size of the micelle).

Our models predict that shell sizes increase with scaffold length, but saturate at minimal and maximal shell sizes in the limit of short or long scaffold molecules. Cargo packaging diminishes with increasing scaffold length (for fixed scaffold-cargo interaction valence), eventually leading to sparsely filled shells. However, increasing the scaffold-cargo valence can restore full shells.

A scaling analysis suggests that microcompartment properties are determined by the relative values of characteristic length scales set by: (i) the scaffold length, (ii) the spontaneous curvature of the protein shell, (iii) shell elasticity, and (iv) excluded volume. Our results show how these characteristic scales could be inferred from experimental measurements of the variation of shell size with scaffold length. Importantly, such measurements could reveal the role of shell spontaneous curvature in driving microcompartment closure, which is currently a key unresolved question.

## I. RESULTS AND DISCUSSION

### A. Computational Results

To simulate scaffold-mediated microcompartment assembly, we extend a model with direct cargo-cargo and cargo-shell pair interactions [45] to include scaffold proteins (section Computational model). Although we keep the model minimal to understand general principles, we are motivated by recent experiments on α-carboxysome assembly [43, 121]. The *α*-carboxysome scaffold, CsoS2, contains three linearly connected flexible domains which include binding sites for the cargo molecule rubisco and the shell hexamer proteins. While it is known that the N-terminal domain contains multiple binding sites for rubisco, as yet there is not strong experimental evidence for which sites interact with hexamers. In many other microcompartment types, the scaffold molecules contain ‘encapsulation peptides’ at their C-terminal domains that interact with hexamers, but these have not been identified for CsoS2. Given the current level of experimental uncertainty, we have started with a simple model for the scaffold: a flexible bead-spring polymer with three domains (section Computational model and Fig. 1). The domain at one end has multiple cargo interaction sites, the domain at the other end binds to shell hexamers, and the middle domain has only repulsive interactions with cargo and shell proteins. In future work we will explore other arrangements of interaction sites on the scaffold. To minimize the number of parameters, we restrict the computational model to a single size of cargo particles and scaffold segments. Thus, the simulations do not account for the small excluded volume size of a CsoS2 segment relative to a rubisco holoenzyme. However, we analyze effects of varying excluded volume using the theoretical model described below.

**FIG. 1.**
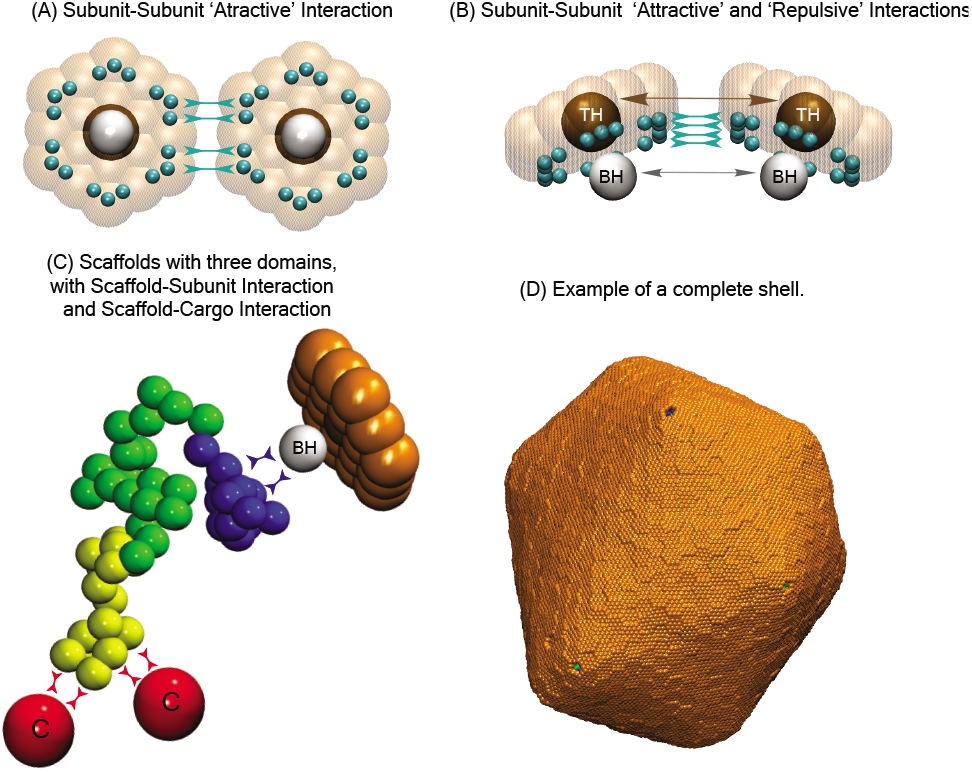
Description of the computational model. **(A)** Bottom and **(B)** side views of shell subunits. Each subunit contains ‘Attractors’ (cyan spheres) on its perimeter, and a ‘Top’ (ocher sphere, ‘T’) and a ‘Bottom’ (white sphere, ‘BH’) in the center above and below the hexamer plane. Interactions between Attractors drive subunit assembly, while Top-Top and Bottom-Bottom repulsions control the subunit-subunit angle and spontaneous curvature radius of the shell *R*_0_. **(C)** Scaffolds are bead-spring polymers with three domains: scaffold-cargo binding domain (yellow) beads have attractive interactions with cargo particles, middle domain (green) beads have no attractive interactions with cargo or shell subunits, and scaffold-shell binding domain (dark blue) beads have attractive interactions with Hexamer Bottom pseudoatoms ‘BH’. The contour length of the scaffold with *N*_s_ beads is denoted as *L*_s_, while the lengths of the domains are *L*_sc_, *L*_sm_, and *L*_sh_, respectively. Excluder atoms (orange pseudoatoms in the plane of the ‘Top’) experience excluded volume interactions with the cargo and scaffold beads. **(C)** Example of a complete shell with spontaneous curvature radius *R*_0_ = 22, with encapsulated cargo and scaffold molecules with *L*_s_ = 64 (*L*_sc_ = 7, *L*_sm_ = 50, *L*_sh_ = 7). All lengths in this article are given in units of *r*_*_, the cargo diameter (the rubisco diameter in carboxysome is ≈13 nm), and energies are in units of the thermal energy *k*_B_*T*.

Based on AFM experiments showing that microcompartment shell facets assemble from pre-formed hexamers [127], our model considers hexamers as the basic assembly units. Carboxysomes assembled in the presence and absence of pentamer proteins have the same structures (except that the latter presumably have vacancies at their vertices) [128]. Thus, to focus on the influence of scaffold molecules on microcompartment size, we employ a minimal model for shell assembly which includes only hexamer proteins. The hexamers have attractive edge-edge interactions with a preferred angle that sets the intrinsic spontaneous curvature radius *R*_0_ of the shell. To further focus on scaffold-mediation, we restrict consideration to purely repulsive direct cargo-cargo and cargo-shell interactions, although attractive cargo-cargo and cargo-shell interactions have not been ruled out in the experimental systems [43, 129]. Moreover, we perform simulations with excess shell proteins to minimize effects of their depletion as assembly proceeds. Additional model details are in the Computational model section.

#### Assembly pathways depend on scaffold-cargo interactions

To elucidate assembly pathways and outcomes, we performed dynamical simulations over a range of scaffold lengths *L*_s_, cargo-binding domain lengths *L*_sc_, shell spontaneous curvature radii *R*_0_, and binding affinities between scaffold-cargo, scaffold-hexamer, and hexamer-hexamer pairs (*ε*_sc_, *ε*_sh_, *ε*_hh_). We find that the strength of scaffold-mediated cargo-cargo interactions strongly influences assembly pathways. Strong interactions (*ε*_sc_ ≳ 1.2 for the concentrations and wild-type-length cargo binding domain length used in our simulations) drive phase separation into a dilute phase and a high density liquid complex of scaffold and cargo molecules. This condition favors two-step assembly pathways, in which the shell proteins adsorb onto the scaffold-cargo domain, and then cooperatively into ordered shell structures. For weaker scaffold-cargo interactions, we observe one-step assembly pathways with simultaneous scaffold-cargo coalescence and shell assembly (Fig. 2 A,B).

**FIG. 2.**
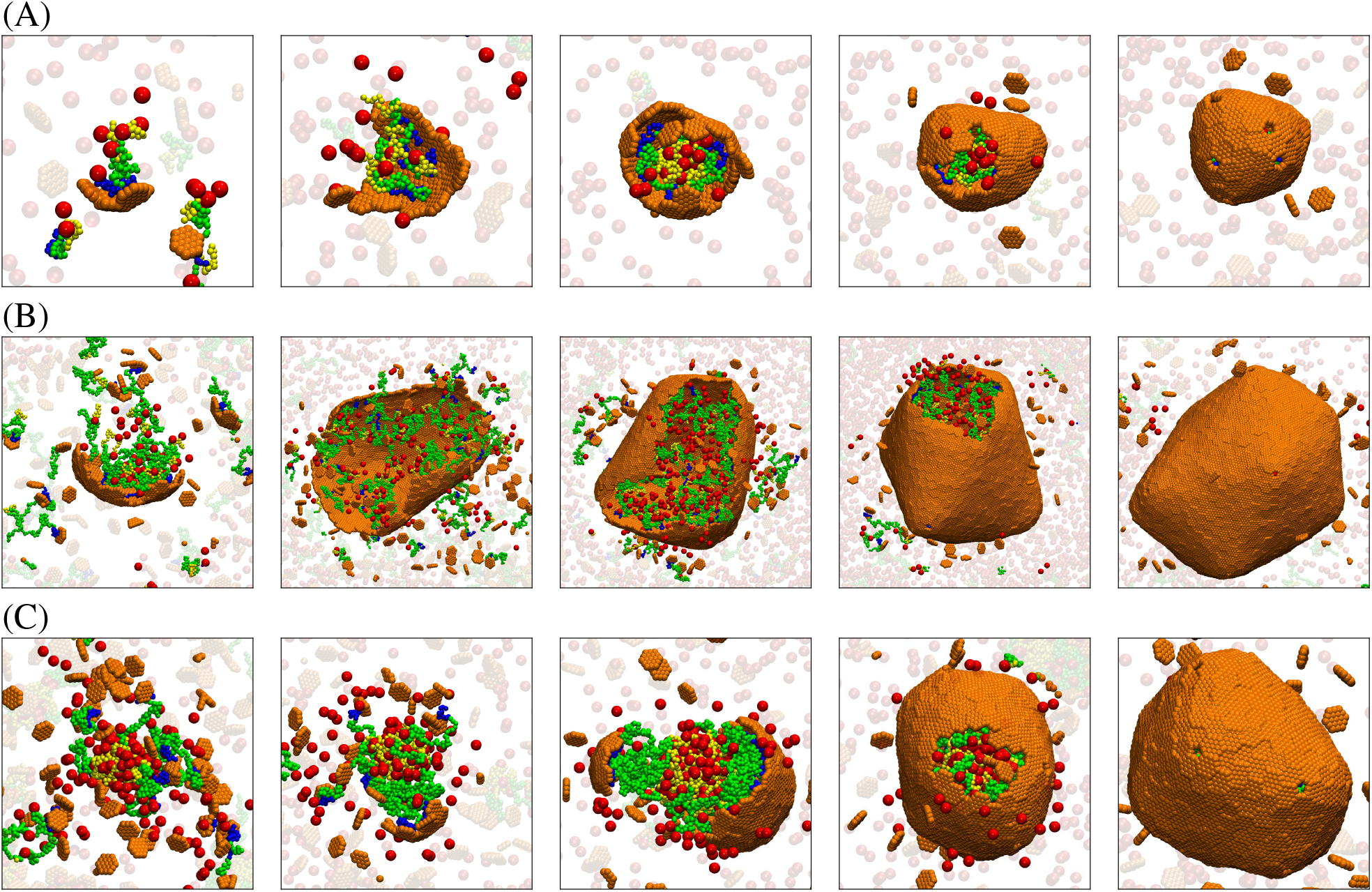
Snapshots from simulation trajectories illustrating typical assembly pathways. **(A)** One-step assembly pathway: Coalescence of the cargo and scaffold occurs concomitantly with shell assembly. The mean size of assembled shells (defined as the radius of gyration of shell subunits) is *R* = 6.3 ± 0.5. Parameter values are: shell spontaneous curvature radius *R*_0_ = 8.0, scaffold contour length *L*_s_ = 24 (*L*_sm_ = 10, *L*_sc_ = 7), and hexamer-hexamer interaction strength *ε*_hh_ = 2.85 (all lengths and energies are given in units of *r*_*_ and *k*_B_*T* respectively). **(B)** One-step assembly, with mean shell size *R* = 18 ± 3.6, for *R*_0_ = 22, *L*_s_ = 64 (*L*_sm_ = 50, *L*_sc_ = 7), and *ε*_hh_ = 2.65. **(C)** Two-step assembly pathway: The scaffold and cargo phase separate prior to shell assembly, due to increased valence of the scaffold-cargo binding domain *f*_sc_ = 0.42. The shell size is *R* = 10.66 ± 0.60. Parameter values are: *R*_0_ = 22, *L*_s_ = 64 (*L*_sm_ = 30, *L*_sc_ = 27), and *ε*_hh_ = 2.65. For (A)-(C), the scaffold-shell and scaffold-cargo interaction strengths are *ε*_sh_ = 2.5 and *ε*_sc_ = 1.0. Snapshots are shown at different scales for clarity.

Here we are motivated by *α*–carboxysomes, for which experiments observe one-step pathways and that the rubisco does not undergo phase separation in the absence of shell proteins at physiological conditions [41, 67, 130]. Therefore, for all further results we set *ε*_sc_ = 1.0, which drives one-step assembly pathways for the wild-type scaffold and cargo-binding domain lengths.

Examples of typical assembly trajectories are shown for scaffolds with: all domains at wild-type lengths, a long middle domain, and a long cargo binding domain in Fig. 2A,B,C respectively. For the first two examples, assembly begins with a small aggregate involving all of the microcompartment constituents: shell subunits, scaffold, and cargo. Because we focus on weak binding affinities, all of these interactions are required to stabilize small aggregates. Consequently, the shell grows concomitant with further coalescence of scaffold and cargo, until shell closure terminates assembly. While the scaffold and cargo are essential for nucleating assembly in both cases, the scaffolds with longer middle domains (which do not attract cargo particles in the model) lead to a lower density of packaged scaffold and cargo (Fig. 2B) compared to the shorter middle domains (Fig. 2A). In contrast, the higher valence cargo-binding domain (Fig. 2C) mediates cargo-cargo attractions that are nearly strong enough to drive phase separation, resulting in pathways closer to two-step assembly and full shells. This result is consistent with experimental observations that *α*-carboxysome scaffold and rubisco can phase separate without shell proteins under non-physiological conditions [37, 43].

#### Shell size depends on scaffold length

We find that shell size depends most sensitively on scaffold length and shell spontaneous curvature. Changing the cargo-binding domain length (at fixed total scaffold length) or binding affinities has only a small effect on shell size (Figs. S3C and S5). Motivated by the experiments, we vary the scaffold length above and below its wild-type length, with *L*_s_ = 24 and *L*_sc_ = 7 model segments roughly corresponding to the native values for the total scaffold length and cargo binding domain length (see SI section S2. Computational Model Details).

Fig. 3A shows the mean size of assembled shells as a function of the middle domain length *L*_sm_ (with two other domains at constant lengths *L*_sc_ = 7, *L*_sh_ = 7) for several values of *R*_0_. We identify two regimes, depending on whether the preferred size of the scaffold molecules (i.e. their end-to-end distance *R*_scaf_ in the absence of shell proteins, (Fig. S2)) is smaller or larger than the intrinsically preferred shell size *R*_0_. For small scaffold lengths, *R*_scaf_ < *R*_0_. In this limit, the number of encapsulated scaffold molecules (*n*_scaf_) relative to shell subunits (*n*_shell_), given by *σ*_s_ = *n*_scaf_/*n*_shell_, decreases with increasing scaffold length (Fig. 3B). Finally, for *R*_scaf_ ≫ *R*_0_ shells are unable to close around scaffold molecules, leading to incomplete assembly.

**FIG. 3.**
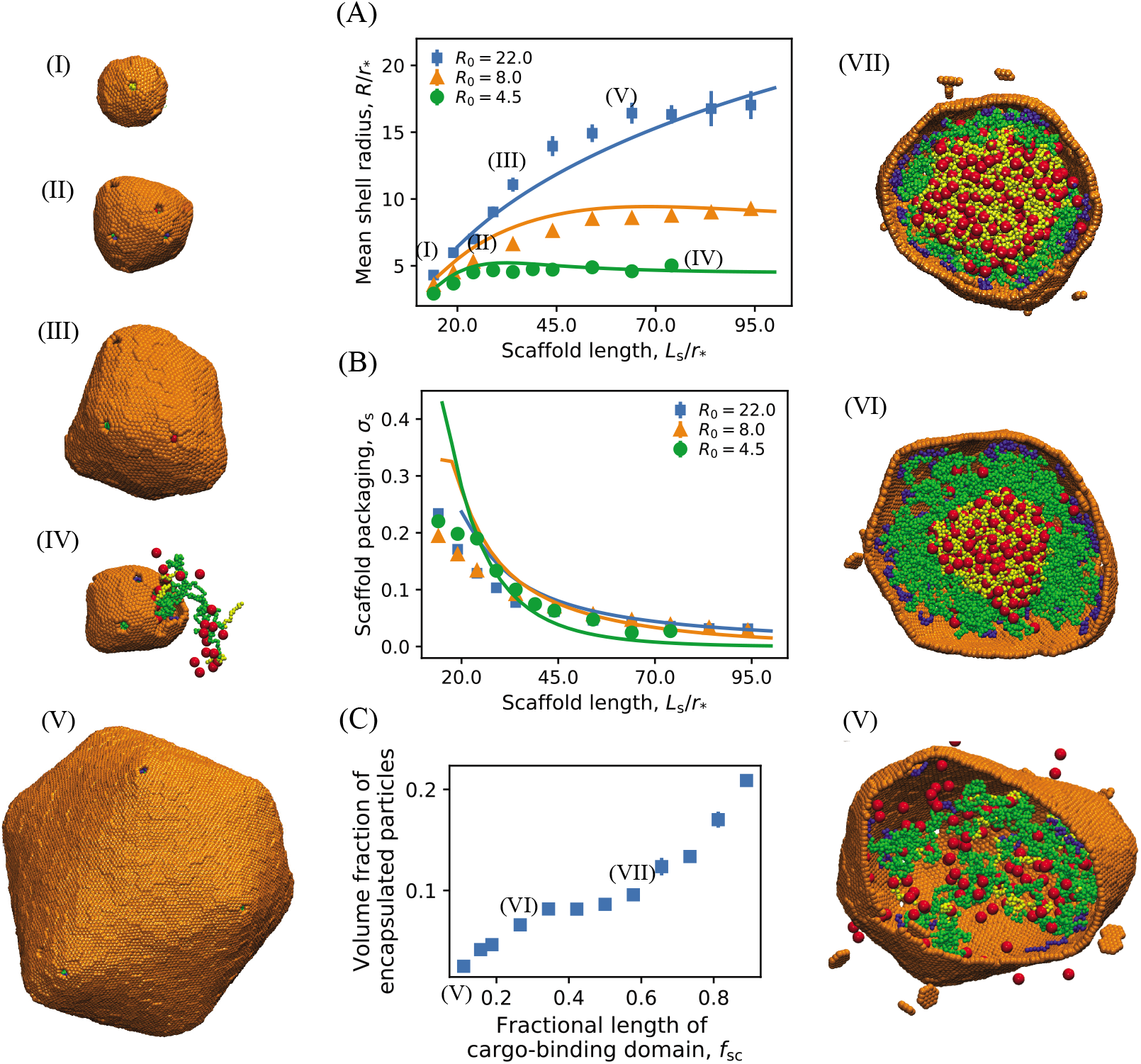
**(A)** The mean shell radius as a function of the scaffold length calculated from Brownian dynamics simulations (symbols) and the equilibrium theory (Eq. 1, lines). Results are shown for different shell spontaneous curvature radii: 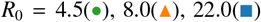. Example snapshots of assembled shells are shown to the left, taken from simulations with (I): *R*_0_ = 22, *L*_s_ = 14, (II): *R*_0_ = 8.0, *L*_s_ = 24, (III): *R*_0_ = 22, *L*_s_ = 34, and (IV): *R*_0_ = 4.5, *L*_s_ = 84, (V): *R*_0_ = 22, *L*_s_ = 64. **(B)** Dependence of scaffold packaging on scaffold length and shell spontaneous curvature. The scaffold surface density (or ratio of packaged scaffold molecules to shell hexamer subunits, *σ*_s_ ≡ *n*_scaf_/*n*_shell_) is shown for the same parameter values as in (A). In (A) and (B) the length of the middle domain of the scaffolds *L*_sm_ is varied, with fixed shell-interacting and cargo-interacting domain lengths *L*_sh_ = 7 and *L*_sc_ = 7. The lines correspond to numerical minimization of Eq. 1, with respect to 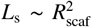 and *σ*, with the scaffold chemical potential as an adjustable parameter set to Δ*μ* = −2.5*k*_B_*T*. Other theory parameters are taken from the simulations: 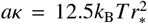 (approximately determined in Ref. 45) and scaffold bead excluded volume 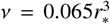. **(C)** Dependence of the volume fraction of encapsulated particles (cargo and scaffolds) on the fractional length of cargo-binding domain of scaffolds *f*_sc_ = *L*_sc_/*L*_s_, for *R*_0_ = 22, *L*_s_ = 64, and *L*_sh_ = 7. Snapshots of the interior of shells assembled at different parameter values are shown on the right: (V): *f*_sc_ = 0.11, (VI): *f*_sc_ = 0.27, (VII): *f*_sc_ = 0.58. Other parameter values for (A), (B) and (C) are as follows. Shell subunit-subunit affinities: *ε*_hh_ = 3.15 at *R*_0_ = 4.5, *ε*_hh_ = 2.85 at *R*_0_ = 8.0 and *ε*_hh_ = 2.65 at *R*_0_ = 22; scaffold-shell interaction *ε*_sh_ = 2.5, and scaffold-cargo interaction *ε*_sc_ = 1.0.

Although the total amount of packaged cargo increases with scaffold length for small lengths, due to the increasing shell size, the volume fraction of packaged cargo and scaffolds decreases monotonically with scaffold length (Fig. S3A and B). I.e., long scaffold middle domains lead to poor cargo loading, because the additional scaffold segments do not attract cargo but increase excluded volume in our model.

#### Longer cargo binding domains increase cargo loading

Fig. 3C shows how cargo packaging depends on increasing the fractional length of the cargo binding domain, *f*_sc_ = *L*_sc_/*L*_s_; i.e., increasing *L*_sc_ at fixed *L*_s_. We see that packaging increases monotonically with *f*_sc_, but there is an inflection point at *f*_sc_ = 0.3, beyond which shells are nearly full. This point corresponds to the cargo binding domain length beyond which phase separation and two-step assembly pathways occur, suggesting that both thermodynamic and kinetic factors can influence cargo packaging.

#### Effect of binding affinity

Finally, we note that the shell size is relatively insensitive to the hexamer-hexamer *ε*_hh_ and scaffold-hexamer *ε*_sh_ affinities, provided they are strong enough to drive assembly and scaffold packaging (Fig. S5). However, the scaffold-cargo complex mediates effective hexamer-hexamer interactions, and thus larger values of *ε*_hh_ are required for complete shell assembly in the large *L*_s_ limit where few scaffold molecules are packaged.

### B. Equilibrium theory

#### Model

To further understand the interplay between scaffold and shell properties, we developed a simple equilibrium model (see SI section S1. Equilibrium Theory). Although kinetic factors may be important as noted above, the thermodynamic model captures key trends observed in the dynamical simulations.

We make the following simplifying assumptions; extensions to eliminate these approximations are straightforward. First, we restrict the theory to spherical shells as observed in the simulations. Second, cargo molecules are implicitly represented by a net driving force for scaffold packaging Δμ that accounts for scaffold-shell and scaffold-cargo attractive interactions, as well as the mixing entropy penalty for scaffold coalescence. We further assume that the coalesced cargo and scaffold-cargo binding domains are concentrated in the shell interior, while the other end of the scaffold, the scaffold-shell binding domain, binds to the inner shell surface. Our model system can thus be thought of as a diblock copolymer spherical micelle (e.g. [122–124, 131–133]), except that the stoichiometry of the interior/exterior block is annealed, and there is an exterior shell with a preferred curvature that may be incommensurate with the optimal size of the micelle within. We discuss below the implications of model predictions for alternative scaffold packaging geometries.

The free energy (per shell subunit) of a shell with radius R and ratio of scaffold/shell subunits *σ*_s_ is given by

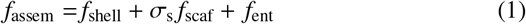

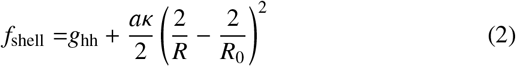

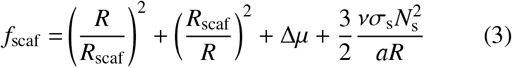

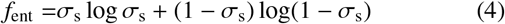

Here *f*_shell_ gives the shell free energy, with the first term (*g*_hh_) the free energy per hexamer due to hexamer-hexamer contacts (see SI section S1. Equilibrium Theory, Eq. S8), and the second term describing the bending energy due to deviations from the shell spontaneous curvature, with *a* the subunit area and *κ* the bending modulus. For simplicity, we neglect contributions from the 12 five-fold defects that are required by topology (in physical and computational microcompartments). Including these will not qualitatively change the results (see SI section S1. Equilibrium Theory and Ref. 45). Then the first two terms in *f*_scaf_ give the entropic penalty for stretching or compressing scaffold molecules from Rscaf. The following two terms account for attractive scaffold-cargo and scaffold-shell interactions (Δ*μ*), and excluded volume interactions, with *v* as the scaffold segment excluded volume. Finally, *f*_ent_ represents the mixing entropy of encapsulated scaffolds. The equilibrium shell size and scaffold loading are obtained by minimizing the total free energy of the system. In the thermodynamic limit, the distribution of shell sizes will be strongly peaked around the size that minimizes the free energy **per subunit** of the shell complex (see SI section S1. Equilibrium Theory and Refs. [45, 134]).

#### Comparison of theory with computational results

The solid lines in Fig. 3A,B show the theory results, with the parameters 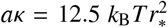 and 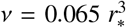 calculated from simulations and Δ*μ* as a fitting parameter chosen by eye. We set 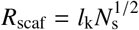 since the theory separately accounts for chain stretching and excluded volume (as in the Flory approximation [135]).

We find that the theory qualitatively captures the main features observed in the simulation results. In particular, the shell size increases with scaffold length, until saturating near *R*_0_ for long scaffolds, where excluded volume limits scaffold packaging. However, for simulations with *R*_0_ =22, longer scaffolds undergo limited coalescence in the center of the shell, causing the shell size to saturate at smaller scaffold lengths than predicted by the theory.

#### Scaling analysis, and the small excluded volume limit

Eq. 1 predicts that the system behavior depends on several characteristic length scales. In addition to the shell spontaneous curvature radius *R*_0_ and scaffold size *R*_scaf_, we obtain length scales from the scaffold excluded volume, *R*_exc_ ~ (*N*_s_*ν*)^1/3^, and the shell elastic energy, 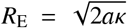. Specifically, *R*_E_ is the radius for which a shell with no spontaneous curvature *R*_0_ = ∞ would have a bending energy of *k*_B_*T* per subunit. Different scaling forms can be deduced from Eq. 1 depending on the relative magnitudes of these lengths.

Since our model is most applicable to biological microcompartments in the limit of small excluded volume (see discussion), we focus here on the limit of small excluded volume, *ν* → 0. In SI section S1. Equilibrium Theory we present a detailed derivation of these expressions and further analysis for finite excluded volume.

Fig. 4 shows the shell size calculated as a function of scaffold length and shell spontaneous curvature with *v* = 0, by minimizing Eq. 1. The dashed black lines encompass the region with significant scaffold packaging for Δ*μ* = −10*k*_B_*T*; the full dependence of scaffold packaging on parameters is shown in Fig. S1. We see that the range of accessible shell sizes depends crucially on the relative sizes of the preferred shell curvature and the elastic length scale. For *R*_0_ > *R*_E_ the shell size is highly tunable with scaffold length, varying over orders of magnitude, while only small deviations from the preferred shell curvature are possible in the opposite limit *R*_0_ < *R*_E_. This predicted trend may be a way to experimentally probe *R*_0_ in microcompartment systems.

**FIG. 4.**
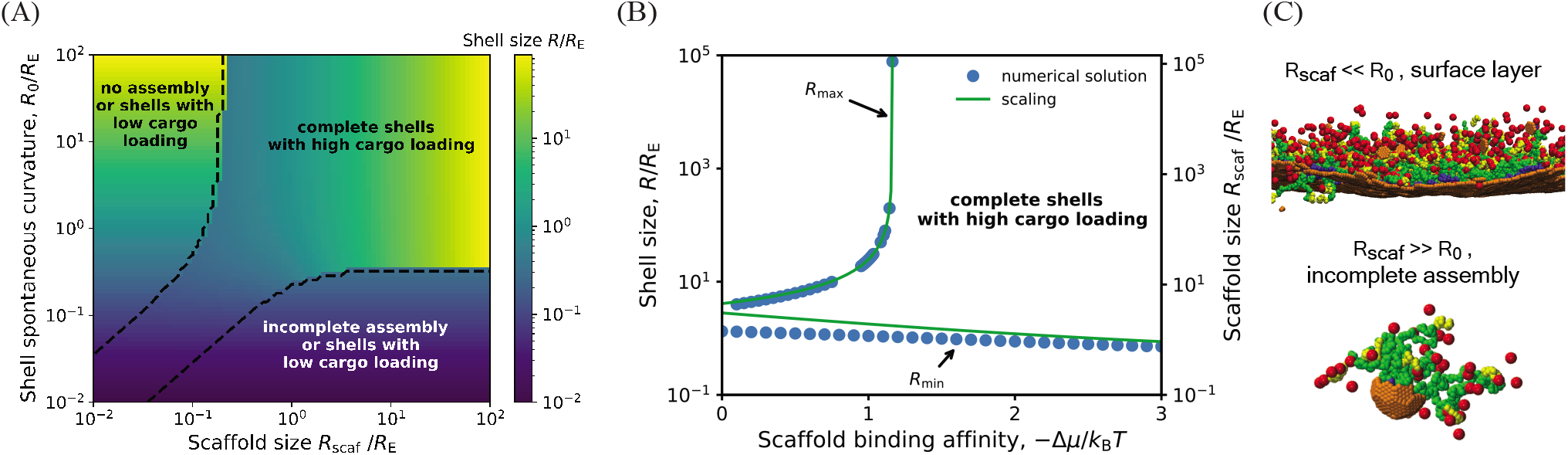
**(A)** Shell radius as a function of the shell spontaneous curvature radius *R*_0_ and scaffold unperturbed size *R*_scaf_ (normalized by the shell elastic energy length scale *R*_E_), calculated by numerically minimizing Eq. 1, with scaffold chemical potential Δ*μ* = −10*k*_B_*T*, *R*_E_ = 30*r*_*_, subunit area 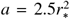, and excluded volume parameter *ν* = 0. The dashed line encompasses parameter values that lead to efficient scaffold packaging (see Fig. S1 for the amount of packaged scaffold as a function of these parameters). Outside of this region *σ*_s_ ≈ 0, leading to assembly of nearly empty shells (with a surface layer of scaffold and cargo) with sizes approximately equal to *R*_0_, incomplete shells, or no assembly. The heat map in this region corresponds to the size of the complete but nearly empty shells. **(B)** The minimum and maximum shell radius (normalized by *R*_E_) predicted by the theory for varying the scaffold length as a function of the scaffold packaging driving force Δ*μ*, for fixed shell spontaneous curvature *R*_0_ = 50. The left y-axis shows the shell size, while the right y-axis shows the corresponding scaffold length, showing that the shell size closely tracks the scaffold preferred size in this regime. The symbols correspond to maximum and minimum shell sizes obtained by numerically minimizing Eq. 1, and the lines show the asymptotic results (Eq. 7). For scaffold lengths above the maximum length, shells would be either incomplete or empty, whereas below the minimum length, shells will either be incomplete or have low packaging of scaffold and cargo. Other parameters are the same as in (A). **(C)** Examples of incomplete assemblies that form in the dynamics simulations with overly short (top) or long (bottom) scaffold molecules. Parameters are (top) *R*_0_ > 300 and *L*_s_ = 34, (bottom) *R*_0_ = 4.5 and *L*_s_ = 84. Other parameters are as in Fig. 3.

##### Scaling analysis

Further insight can be obtained from a simple scaling analysis. First, minimizing the free energy with respect to *σ*_s_ gives

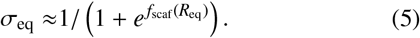

Eq. 5 shows that scaffold/cargo packaging diminishes as the shell size deviates from the preferred scaffold size *R*_scaf_, but that scaffold can be packaged over a greater range of shell sizes by increasing |Δ*μ*|. Second, we obtain simplified expressions for the shell size and packaging depending on the relative characteristic scales as follows.

##### Small intrinsic shell curvature

*R*_0_ ≫ *R*_E_. In this limit the optimal shell size becomes asymptotically independent of shell spontaneous curvature, and minimizing the free energy results in an optimal shell size

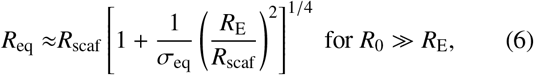

which can be further simplified in two limits:

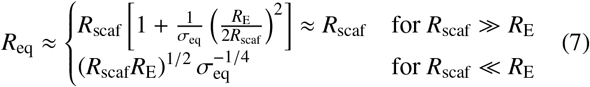

Thus, the shell size depends on a competition between the preferred scaffold size and the tendency of the shell proteins to disfavor curvature: the shell size tracks the scaffold size *R*_eq_ ≈ *R*_scaf_ for long scaffolds, but adopts the geometric mean of the scaffold size and shell elastic length for short scaffolds. These scaling estimates are compared against the full free energy in Fig. 5A.

**FIG. 5.**
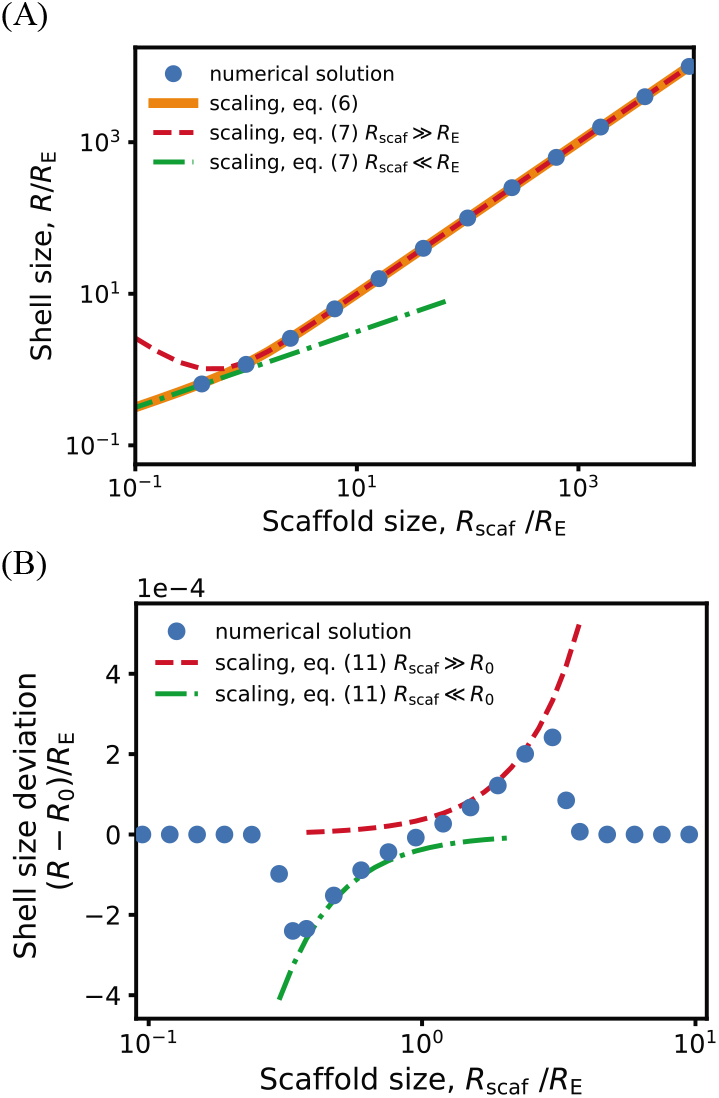
Dependence of shell size on the scaffold preferred size *R*_scaf_ calculated from the asymptotic analysis, (lines, Eqs. (7) and (11), compared against numerical minimization of Eq. 1 (symbols). Results are shown in **(A)** and **(B)** respectively for shell spontaneous curvature radius values that are large or small compared to the shell elastic length scale: **(A)** *R*_0_ = 300 and **(B)** and *R*_0_ = 1, with *R*_E_ = 30. Other parameter values are 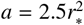, Δ*μ* = −10*k*_B_*T*, and *v* = 0.

However, Eq. 5 shows that scaffold packaging will expo-nentially diminish for large deviations from *R*_scaf_. Therefore, we estimate upper and lower bounds on the scaffold-driven changes in shell size by noting that shell bending energy must be compensated by the free energy of scaffold packaging:

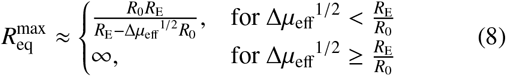

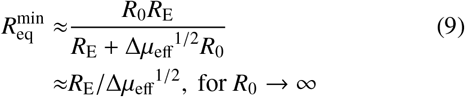

with Δ*μ*_eff_ ≡ −(Δ*μ* + 2*k*_B_*T*)*σ*_eq_ – *f*_ent_ the effective confinement free energy per shell subunit, since *f*_scaf_ ≈ Δ*μ* + 2*k*_B_*T* for *R* ≈ *R*_scaf_, and

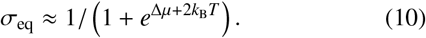

The packageable range of scaffold lengths can then be de-termined by combining Eqs. 8, 9 and 7 (Fig. 4B)). Outside of these bounds, scaffold packaging decreases exponentially, resulting in either no assembly or assembly of empty shells with sizes near *R*_0_. Alternatively, for small scaffolds, if the scaffold-shell energy is sufficient to drive scaffold packaging without cargo, the scaffold can form a layer in the vicinity of the shell surface (Fig. 4C). That scenario is not accounted for in the theoretical model.

Importantly, in any of these scenarios, the microcompartments will saturate at a *minimum* shell size with decreasing scaffold length (Eq. 9), which is an experimentally testable prediction.

##### High intrinsic shell curvature

*R*_0_ < *R*_E_. In this regime the preferred curvature of the shell proteins influences the assembled shell size, so we must consider the full free energy Eq. 1. Then, the limits in which the preferred scaffold size is larger or smaller than *R*_0_ result in

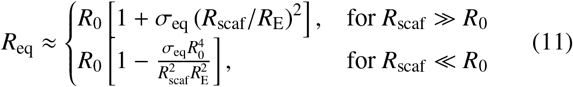

showing that the shell size depends on the competition between the shell spontaneous curvature and scaffold size, with relative importance determined by the shell bending modulus (via *R*_E_). As above, there are bounds on the range of scaffold lengths that can be efficiently packaged, which increase with |Δ*μ*|. Since the shell bending energy dominates in this limit, the shell size exhibits only small deviations from *R*_0_, and the extent of scaffold incorporation is determined by the competition between the packaging driving force Δ*μ* and the scaffold confinement free energy *f*_scaf_, resulting in:

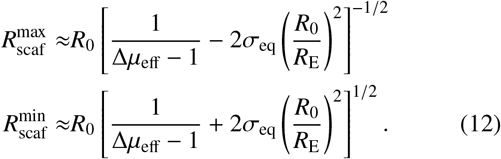

The range of shell sizes 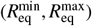 can then be determined by combining Eqs. 10–12. Consistent with the numerical solution (Figs. 4 and S1), the range of accessible shell sizes is considerably smaller than in the large *R*_0_ limit.

## II. CONCLUSIONS

We have theoretically and computationally investigated the equilibrium behavior and dynamical assembly pathways of a system in which polymeric scaffold molecules drive coalescence of many cargo particles, and their packaging within an assembling protein shell. While motivated by biological systems such as bacterial microcompartments, we have employed minimal models allowing for generic conclusions about such multicomponent assemblies. Thus, in addition to elucidating the behaviors of natural and reengineered bacterial microcompartments, our results suggest design principles for tailoring the size, morphology, and loading of synthetic or biomimetic systems such as delivery vehicles.

We find that the shell size and the amount of packaged cargo are determined by several characteristic length scales: the preferred sizes of the protein shell and the scaffold, the scaffold excluded volume size, and a characteristic elastic length scale of the shell (which depends on its rigidity). In our computational model, we have considered a relatively low bending rigidity of the shell (motivated by experimental observations on bacterial microcompartments), and scaffold excluded volume plays a significant role. In this case, results depend on whether the preferred scaffold size is either smaller than or larger than the spontaneous curvature radius. In the limit of small scaffold molecules, assembly driving forces are dominated by stretching of the scaffold between the cargo-filled interior and the surface of the protein shell, leading to shells which are smaller than the spontaneous curvature radius (provided that the scaffold-shell interactions and scaffold-mediated cargo-cargo interactions are stronger than shell bending elasticity). In the opposite limit of long scaffold molecules, scaffold packing becomes a dominant contribution, leading to decreased scaffold-cargo packaging and a shell size that roughly saturates at the spontaneous curvature radius.

Our simulations also demonstrate that the amount and morphology of packaged cargo depends on the strength and geometry of scaffold-cargo interactions. Multivalent scaffold-cargo interactions are essential for achieving full shells, while the scaffold length and distribution of cargo binding sites on the scaffold influence the morphology of the packaged cargo. When cargo-binding sites are limited to one end of long scaffold molecules, with scaffold-shell interactions at the other end, we observe a tri-layer micelle-like complex, with an interior core of cargo, surrounded by a layer of scaffold, and finally the shell on the exterior. In contrast, longer scaffold-cargo binding domains or shorter scaffolds lead to a nearly uniform distribution of cargo and scaffold in the shell interior.

Because our primary objective was to qualitatively understand scaffold-mediated shell assembly, we did not tune the computational model parameters to match a specific biological system, and the computational model likely overestimates the scaffold excluded volume compared to bacterial microcompartments. For example, the *α*-carboxysome scaffold (CsoS2) accounts for only 12% by mass of a native *α*-carboxysome [136], compared to 70% in our model. Moreover, while we have included very weak scaffold intersegment attractions in our model, stronger inter-segment attractions among CsoS2 residues could further reduce the effective excluded volume.

To facilitate more direct comparisons against experiments, we also developed a more tractable, though simplified, equilibrium theoretical model. Using this model, we have explored a wide parameter range, including the effect of reducing the excluded volume. The theoretical analysis suggests that the ability of a scaffold to modulate shell size depends crucially on whether the protein shell spontaneous curvature radius is large or small compared to its elastic length scale. In the regime of a large spontaneous curvature radius, shell size is extremely malleable and can be tuned over a wide range by varying scaffold length. In contrast, shell proteins with a small preferred curvature radius exhibit only small size fluctuations, and relatively poor packaging of scaffold molecules with incommensurate sizes. In both regimes, the range of scaffold lengths that can be packaged effectively, and correspondingly the range of accessible shell radii, increase with the free energy driving force for scaffold incorporation. This important control parameter depends on the scaffold concentration and the valence and affinity of the scaffold-cargo and scaffold-shell domains.

### Testing against experiments

Since our models are general, these predictions could be tested in a variety of biological or synthetic systems. However, bacterial microcompartments are the existing system in which testing would be most straightforward, especially due to the modular nature of microcompartment scaffold molecules. For example, the CsoS2 scaffold of the α-carboxysome can be represented as a tri-block co-polymer, with the cargo-binding domain in the N-terminal block, and shell-binding sites in the C-terminal block. Our results could thus be compared against experiments in which mutagenesis [43] is used to change the length of the middle domain (Fig. 3A,B) or the cargo-binding domain (Fig. 3C), although we note that the presence of shell binding sites in the middle block has not been ruled out.

Importantly, our results suggest a path to assess the magnitude and importance of the spontaneous curvature, a key unresolved question in microcompartment assembly. In particular, recent experiments on recombinant systems exhibit assembly of small (~ 20 - 30 nm) empty microcompartment shells, suggesting a small shell spontaneous curvature radius. Yet, other experiments observe extensive flat sheets of hexamers assembling on surfaces, suggesting at least the possibility of a low or zero spontaneous curvature. In this regard, both the simulations and theoretical model predict that shells with a small spontaneous curvature radius exhibit only narrow fluctuations in size when presented with scaffold in molecules of different lengths, whereas shells with a large spontaneous curvature radius can assemble over a wide range of sizes to accommodate scaffolds with different lengths. In the small spontaneous curvature regime, our results suggest that preferred shell curvature plays a key role in driving shell closure, whereas in the large-curvature radius regime scaffold properties (this work) and cargo properties[45, 46] are important for shell closure.

One readily testable prediction from our model is that there is a minimum accessible shell size. Reducing scaffold lengths below this limit will either lead to a saturating shell size with diminished scaffolding packaging, or abrogation of shell assembly.

### Outlook

Finally, our results suggest additional physical ingredients that will be important to study for understanding biological microcompartments or guiding the design of synthetic delivery vehicles. First, increasing the rigidity of the scaffold molecules would diminish the range of variation of shell sizes. This trend can be seen from the equilibrium scaling of the shell size in Eqs. 7 and 11, since the unperturbed scaffold size increases with its rigidity. Second, combining direct cargo-cargo interactions with scaffold mediation could enhance cargo packaging within assembling shells, since. This proposition is based on our observation in this work that increasing effective scaffold-mediated cargo interactions promotes cargo loading, and is consistent with observations from previous models with cargo coalescence driven only by direct cargo-cargo pair interactions [44]. Third, although we observe qualitative agreement between our dynamical simulation results and the equilibrium theory, kinetic effects likely play important roles in some parameter regimes. For example, we have primarily focused on parameters that lead to one-step assembly pathways as observed for *α*-carboxysomes, but we anticipate that kinetic effects will become more important in the case of two-step assembly pathways. Thus, models such as developed here, combined with experiments on different microcompartment systems and solution conditions, are needed to fully understand the interplay between thermodynamics and kinetics in scaffold-mediated shell assembly and cargo packaging.

## III. METHODS

### A. Computational model

Our computational model extends a previous model for assembly of a shell around fluid cargo[44, 45] to include scaffold molecules and scaffold-mediated cargo-cargo and cargo-shell interactions. For simplicity, we do not include direct cargo-cargo and cargo-shell attractive interactions, and we consider only hexamer shell proteins. The interactions among components are as follows, with further details given in SI section S2. Computational Model Details.

#### Scaffolds

In carboxysome systems, attractions between rubisco particles are mediated by auxiliary proteins (*e.g*. the intrinsicallly disordered protein CsoS2 in *α*–carboxysomes [41, 43] and the protein CcmM in *β*–carboxysomes [38]). The *α*–carboxysome scaffold molecule, CsoS2, contains three linearly connected flexible domains: the C-terminal domain contains multiple binding sites for the cargo molecule rubisco, while there is not yet clear experimental evidence for the location of the shell binding sequences. However, in *β*-carboxysomes and other bacterial microcompartments, shell-cargo attractions are mediated by an ‘encapsulation peptide’ sequence in the N-terminal domain of the scaffold [23, 31, 38, 137, 138]. Motivated by these characteristics, we model the scaffold as a flexible bead-spring polymer with three linearly connected domains: a scaffold-cargo binding domain at one end with multiple cargo interaction sites, a scaffold-shell binding domain at the other end with multiple shell hexamer binding sites, and a middle domain that has only repulsive interactions with cargo and shell proteins. Based on experimental evidence, we include weak attractive interactions between pairs of scaffold beads.

The interactions between various molecule types are modeled as follows.

#### Scaffold-cargo interactions

Attractive interactions between cargo particles and the ‘cargo-interacting’ beads (type ‘SC’) in scaffolds are modeled by a Lennard-Jones potential with well-depth parameter *ε*_sc_.

#### Scaffold-shell interactions

Attractive interactions between hexamers and scaffolds are modeled by a Morse potential with well-depth parameter *ε*_sc_, between beads in ‘shell interacting’ beads in polymers (blue beads in Fig. 1C) and Bottom pseudoatoms on hexamers (type ‘BH’). We also add a layer of ‘Excluders’ in the plane of the ‘Top’ pseudoatoms, which account for shell-scaffold and shell-cargo excluded volume interactions.

#### Shell-shell interactions

Interactions between edges of BMC shell proteins are primarily driven by shape complementarity and hydrophobic interactions [51]. To mimic these short-ranged directionally specific interactions, each model subunit contains ‘Attractors’ on its perimeter that mediate shell-shell attractions. Complementary Attractors on nearby subunits have short-range interactions (modeled by a Morse potential, Eq. S26 in SI section S2. Computational Model Details). Attractors which are not complementary do not interact. The arrangement of Attractors on subunit edges is shown in Fig. 1, with pairs of complementary Attractors indicated by cyan double-headed arrows. The shell-shell binding affinity is proportional to the well-depth of the Morse potential between complementary Attractors, given by *ε*_hh_.

To control the shell spontaneous curvature and bending modulus, each subunit contains a ‘Top’ (type ‘TH’) pseudoatom above the plane of Attractors, and a ‘Bottom’ pseudoatom (type ‘BH’) below the Attractor plane. There are repulsive interactions (WCA interactions, Eq. S25) between Top-Top, Bottom-Bottom, and Top-Bottom pairs of pseudoatoms on nearby subunits. The relative sizes of the Top and Bottom pseudoatoms set the preferred subunit-subunit binding angle (and thus the spontaneous curvature *R*_0_), while the interaction strength (controlled by the well-depth parameter *ε*_angle_) controls the shell bending modulus *κ*. The Top-Bottom interaction ensures that subunits do not bind in inverted orientations [77]. Since the shell-shell interaction geometries are already controlled by the Attractor, Top, and Bottom pseudoatoms, we do not consider Excluder-Excluder interactions.

### B. Simulations

We simulated assembly dynamics using the Langevin dynamics algorithm in HOOMD (which uses GPUs to efficiently simulate dynamics [139]), and periodic boundary conditions to represent a bulk system. The subunits were modeled as rigid bodies [140]. Each simulation was performed in the NVT ensemble, using a set of fundamental units [141] with the unit length scale *r*_*_ defined as the cargo particle diameter, and energies in units of the thermal energy, *k*_B_*T*. The simulation time step was 0.005 in dimensionless time units, and we performed 2 × 10^8^ timesteps in each simulation, except the maximum simulation time was increased to 4 × 10^8^ for *L*_s_ >= 40 and *R*_0_ = 22 (because longer timescales were required for the large shells that assemble at those parameters).

#### Systems

Each system included 2300 hexamers, 2000 cargo particles, and 213 scaffold molecules, initialized randomly on a grid in a cubic box with side length 120*r*_*_. For systems that are near the threshold for cargo-scaffold phase separation in the absence of shell subunits because of long scaffold-cargo binding domains (*L*_sc_ > 7), we eliminated correlations imposed by the grid by performing an initial dynamics with excluded volume interactions only (all attractive interactions turned off) for 5 × 10^6^ timesteps. The attractive interactions were turned on after completion of this initialization dynamics.

#### Sample sizes

We performed a minimum of 10 independent trials at each parameter set. Simulations with a large spontaneous curvature radius and long scaffolds, *R*_0_ = 22 and *L*_s_ > 40, result in relatively few complete shells per simulation. Therefore, for these parameters we performed additional trials such that at least 10 complete shells formed amongst all trials at a given parameter set.

#### Connecting simulation nondimensional units to physical values

Although the model is designed to be generic, we are particularly motivated by α-carboxysomes. We can *approximately* map our computational model to carboxysomes by setting the cargo diameter (the unit length scale in the model) to that of the rubisco holoenzyme, implying *r*_*_ ≈ 13 nm. However, to enable tractable simulation of long assembly timescales, we have set the size ratios of hexamer/cargo and scaffold bead/cargo to be larger than the ratios of these proteins relative to rubisco. In particular, our model hexamers have a side length of *r*_*_ and are thus about three times larger than carboxysome hexamers (side-length ≈ 4 nm). To represent the carboxysome system as closely as possible despite this approximation, we have set the bending modulus of the computational model to obtain a characteristic elastic length scale 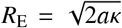 similar to that of carboxysomes. Nanoindentation measurements on *β*-carboxysome shells obtained *κ* ≈ 25*k*_B_*T* [142] (smaller than that for a typical viral capsid), resulting in an elastic length scale of *R*_E_ ≈ 47 nm. Correspondingly, we set the computational bending modulus to about 5 - 10*k*_B_*T* (see SI section S2. Computational Model Details), and thus *R*_E_ ≈ 38 – 54 nm (using a subunit area 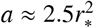).

We perform a similar approximate mapping for the model scaffold molecule, by attempting to match its end-to-end vector (when unconfined) to that of a CsoS2 molecule. The model scaffold behaves as a self-avoiding polymer with a statistical segment length of about one segment, or *l*_k_ ≈ 0.5*r*_*_, and thus has a end-to-end vector 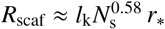, with *N*_s_ = *L*_s_/*l*_k_ (see Fig. S2), with a proportional radius of gyration. Since most of the CsoS2 sequence is thought to be intrinsically disordered, we assume that it also behaves as a self-avoiding polymer. The Halothiobacillus neapolitanus CsoS2 molecule studied in mutagenesis experiments has 869 amino acids. While its radius of gyration has not been reported, the Prochlorococcus CsoS2 with a similar sequence length (792 amino acids) has a radius of gyration of ~ 70 nm [41]. To match this value, we assign the wild-type length of the model scaffold to be *L*_s_ = 25*r*_*_ (50 beads).

## ACKNOWLEDGMENTS

We are grateful to Luke Oltrogge and David Savage for insightful discussions. This work was supported by Award Number R01GM108021 from the National Institute Of General Medical Sciences and the Brandeis Center for Bioinspired Soft Materials, an NSF MRSEC, DMR-1420382 and DMR-2011846. Computational resources were provided by NSF XSEDE computing resources (XStream, Bridges, and Comet) and the Brandeis HPCC which is partially supported by DMR-1420382.

## SUPPLEMENTARY INFORMATION

### S1. EQUILIBRIUM THEORY

#### Free energy

We consider shells composed of *n*_shell_ = 4*πR*^2^/*a* hexamer subunits, with *R* the shell radius and *a* the subunit area, and *n*_scaf_ scaffolds. We incorporate the presence of cargo implicitly through effective attractions between scaffold molecules embodied in the parameter Δ*μ*. The net free energy of an assembling shell with radius *R* is given by

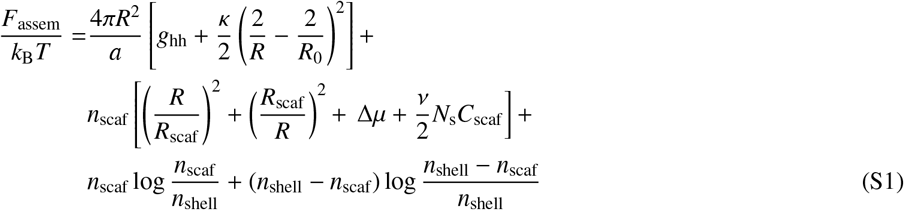

The first term on the right-hand side of Eq. S1 represents the free energy due to subunit-subunit contacts that drive shell assembly, with *g*_hh_ per energy per subunit. The second term describes the elastic energy associated with deviations of the shell from its intrinsic spontaneous curvature, with *κ* the bending modulus. The next two terms give the entropic penalty for stretching or compressing scaffold molecules from their preferred length, *R*_scaf_. In this model we have assumed that the scaffold bridges from the cargo in the interior to the shell surface and thus we set the scaffold length equal to the shell radius. The following two terms account for attractive scaffold-cargo and scaffold-shell interactions, and scaffold-scaffold excluded volume interactions, with the excluded volume per scaffold segment given by *ν* and the scaffold concentration in the shell given by 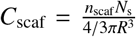. The final two terms represent the mixing entropy of *n*_scaf_ scaffolds binding to a shell with *n*_shell_ subunits. For simplicity we have assumed that one scaffold can bind to each shell subunit, so the total number of binding sites is equal to *n*_shell_.

By defining the surface density of the scaffolds 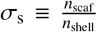, the total free energy per subunit 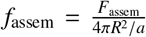 of an assembled shell with radius *R* can be written as

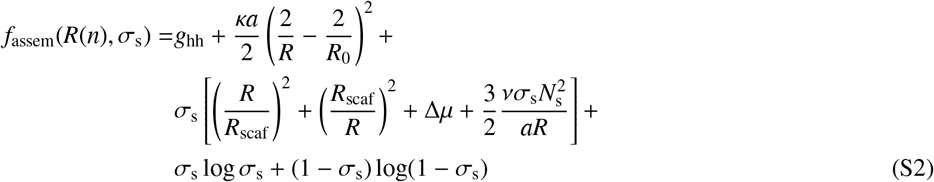

#### Optimal shell size

For all of the theoretical results presented in this work, we consider the case of limiting hexamer subunits, with the scaffold present in excess at fixed chemical potential *μ*_scaf_, so that the term Δ*μ* in Eq. S2 is a constant. The equilibrium distribution *ρ*(*n, σ*_s_) of shells with *n* hexamer subunits can then be calculated by variationally minimizing the total system free energy density

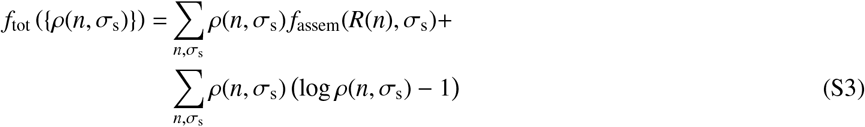

under the constraint of mass conservation: 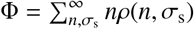 with Φ the total concentrations of hexamer subunits. This obtains a law of mass action

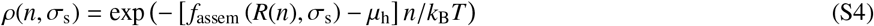

with *f*_assem_ given by Eq. S2 and *μ*_h_ = *k*_B_ *T* log(*ρ*_h_ *ν*_0_) the chemical potentials of free hexamer subunits at their equilibrium concentration *ρ*_h_ with *ν*_0_ a standard state volume. The optimal shell size *n*\* is then obtained by variationally minimizing Eq. S4 with respect to *n* and *σ_s_*. The first of these operations results in

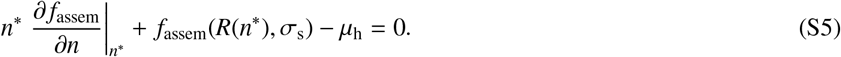

It can be shown that above the pseudo-critical subunit concentration (i.e. the hexamer concentration above which there is significant assembly), the term *f*_assem_(*n*\*, *σ*_s_) – *μ*_h_ ≈ 0 in the limit *n*\* ≫ 1 [134]. To see this, note that at equilibrium the chemical potential of free subunits *μ*_h_ must be equal to the chemical potential of a subunit in a shell, given by

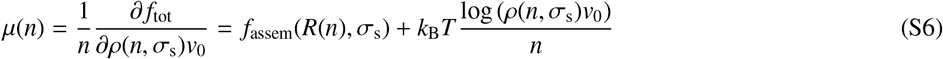

with the last term in Eq. S6 accounting for the per-subunit free energy due to shell mixing entropy. In the limit of large shell size, the last term becomes negligible for shells at the highest concentrations (i.e. with *n* = *n*\*), resulting in *f*_assem_(*R*(*n*\*), *σ*_s_) ≈ *μ*_h_.

Thus, the peak in the shell size distribution corresponds to the minimum free energy **per subunit**; i.e., 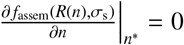, or alternatively:

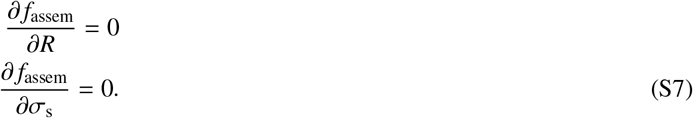

In Ref. 134 this calculation is performed rigorously, showing that for finite concentration the optimum shifts to smaller shell sizes, but that this shift is negligible for large *n*\*.

Finally, note that an analogous analysis could be performed for limiting scaffold molecules, in which case the peak assembly size would correspond to the minimum free energy per scaffold molecule.

Binding free energy of shell subunits. The per subunit shell binding free energy *g*hh can be written as

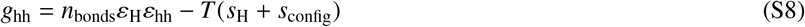

with *n*_bonds_ = 3 the number of bonds per subunit in the shell, *s*_hex_ the translational and rotational entropy penalty associated with binding of hexameric subunits and *s*_config_ = −log(6) accounting for the configurational entropy associated with the subunit’s hexameric symmetry. *ε*_H_ = −3.15 and *s*_H_ = −17.7 are taken from Ref. 44, where they were measured from the dimerization equilibrium constant in simulations of subunits of a similar model only capable of forming dimers.

Approximations in the elastic energy. For simplicity we have modeled shell bending energy using the Helfrich energy for a fluid membrane. We note that for an elastic shell tiled by hexamer subunits, there are additional contributions arising from energy associated with the 12 five-fold disclinations subunits required by topology, and elastic interactions between these disclinations. However, as shown in Ref. 45, the free energy from the defects themselves simply renormalizes the intrinsic spontaneous curvature radius of the shell, with a favorable or unfavorable defect free energy respectively decreasing or increasing the effective spontaneous curvature radius. Since this does not qualitatively change the results, we have neglected this term for simplicity here. We note that while the defect energy in an elastic shell is typically positive (unfavorable), in the case of microcompartments it may be positive or negative since the disclinations are filled by pentamer subunits, which have a different protein sequence (and thus potentially different binding affinity and different preferred binding angles) from the hexamer and pseudo-hexamer proteins that tile the rest of the shell.

The nature of the inter-defect interactions depends on the dimensionless Fopl von Karmann number (*γ* ≡ *YR*^2^/*κ* for a shell of radius R with Young’s modulus Y and bending modulus *κ*). Below a critical value of the Fopl von Karmann number (*γ* = 154), the shells remain roughly spherical [143, 144] and the energy of the inter-disclination elastic interactions is proportional to the shell area. Thus, its contribution can be subsumed into the hexamer-hexamer interaction *g*_hh_ (whose contribution also is linear in shell area) [45]. Above the critical Fopl von Karmann number the shells facet and the inter-disclination energy is screened [143]. Thus, the form of the Helfrich energy is sufficient to describe the equilibrium bending energy of the spherical shells that we consider. However, we note that the inter-defect interactions can have a significant effect on the energy landscape and preferred geometry of assembly intermediates [104, 105], and thus would be important to consider for assembly pathways and kinetics.

### C. Scaling analysis

Here we present a more detailed derivation of the asymptotic results presented in the main text.

#### 1. Optimal shell size

##### A. Low intrinsic shell curvature, *R*_0_ ≳ *R*_E_

In this limit the contribution of the intrinsic shell curvature can be neglected, so the shell and scaffold free energy terms in Eq. 1 of the main text simplify to

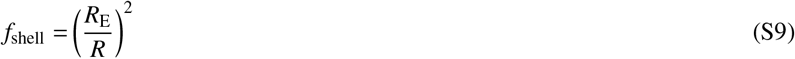

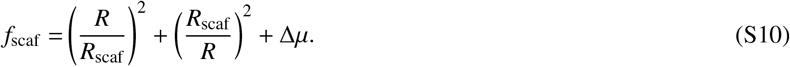

Minimizing the free energy at fixed scaffold incorporation σ then results in an optimal shell size given by Eq. 6 of the main text.

##### B. High intrinsic shell curvature, *R*_0_ < *R*_E_

In this regime the preferred curvature of the shell proteins influences the assembled shell size, so we must consider the full free energy Eq. S2 except we retain the approximation *ν* = 0. As above, we consider separately the limits in which the preferred scaffold size is larger or smaller than the characteristic shell size.

1. *R*_scaf_ ≫ *R*_0_: In this case we simplify the scaffold confinement free energy by neglecting the stretching term:

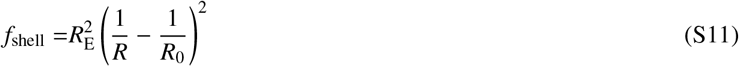

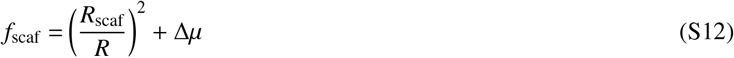

and minimizing the free energy results in

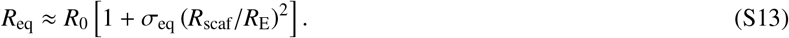
2. *R*_scaf_ ≪ *R*_0_: In this limit we retain the scaffold stretching free energy term:

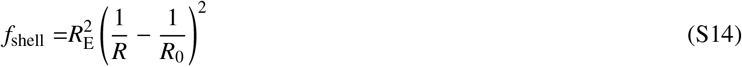

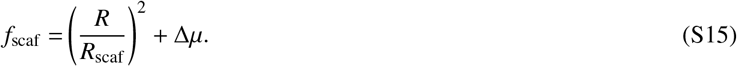

Recognizing that the shell size will remain close to the shell protein spontaneous curvature radius, we set

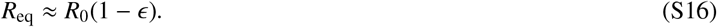

Inserting this into Eq. S16, minimizing with respect to *ϵ* and *σ*, and expanding to linear order in *ϵ* results in

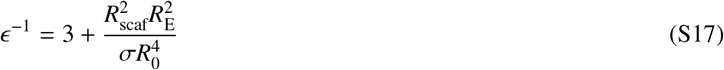

and an optimal shell size

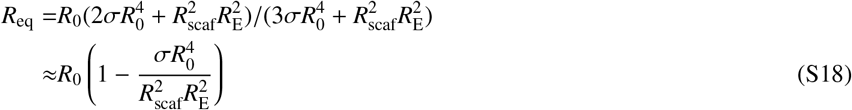

#### 2. Maximum and minimum optimal shell sizes

Here we present details on estimating the maximum and minimum shell sizes that lead to significant scaffold packaging.

##### A. Small intrinsic shell curvature, *R*_0_ ≳ *R*_E_

In this limit the scaffold confinement free energy dominates over shell bending energy, and thus we assume the confinement free energy is approximately given by its minimum value *f*_scaf_ ≈ Δ*μ* + 2*k*_B_*T*. Scaffold packaging is then determined by a competition between the free energy driving scaffold packaging Δ*μ* and the shell bending energy:

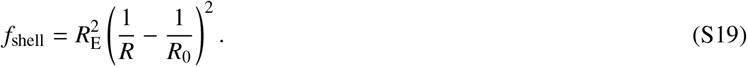

Including the scaffold confinement and mixing free energy terms results in an effective free energy per shell subunit given by:

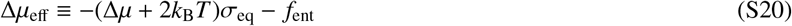

Setting Δ*μ*_eff_ = *f*_shell_ results in two solutions corresponding to Eqs. (8) and (9) of the main text.

##### B. High intrinsic shell curvature, *R*_0_ < *R*_E_

As shown above, the shell size remains close to *R*_0_ in this limit, so scaffold packaging depends on a competition between its confinement free energy *f*_scaf_ and its attractive scaffold-cargo and scaffold-shell interactions represented by Δ*μ*. The nature of this competition depends on whether the preferred scaffold size is larger or smaller than the shell size as follows.

1. *R*_scaf_ > *R*_0_. In this case we focus on the scaffold confinement free energy term, and balance this against Δ*μ*:

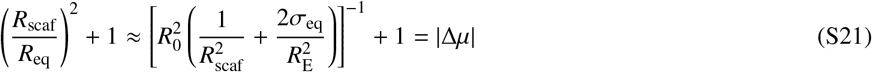

where the +1 on the left hand side of Eq. S21 accounts for the scaffold stretching term. Rearranging to solve for *R*_scaf_ results in the maximum shell size 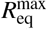 given in Eq. 12 of the main text.
2. *R*_scaf_ < *R*_0_. Here we balance the scaffold stretching term against Δ*μ*:

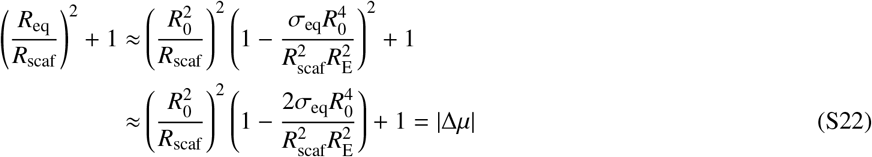 Solving for *R*_scaf_ and taking the limit *R*_0_ ≪ *R*_E_ results in 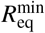 given in Eq. 12 of the main text.

### S2. COMPUTATIONAL MODEL DETAILS

Our model represents subunits as rigid bodies comprised of pseudoatoms arranged to capture the directional attractions and shape of microcompartment pentamer and hexamer oligomers. In comparison to earlier studies with patchy spheres (e.g. [89–92]), multi-pseudoatom subunits better describe the subunit excluded volume shape [74,145,146], which we find to be important for representing assembly around many-molecule cargoes. See Ref. 81 for a comparison of these approaches.

#### Scaffold molecules

Scaffolds are modeled as flexible bead-spring polymers with three domains; cargo-binding beads are at one end, shell-binding beads are at the other end, and beads in the middle domain have only repulsive interactions with cargo molecules and shell subunits. There are weak attractive interactions between pairs of scaffold beads.

The attractive scaffold-cargo interactions lead to collapse of the scaffold-cargo domain, resulting in a shorter effective length of encapsulated scaffolds relative to their free end-to-end size. Therefore, to compare simulation results against the theoretical model (in Fig. 3)), we estimated the effective length of the scaffold-cargo domain that would result in a comparable size, resulting in an effective length of 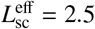 for the actual value *L*_sc_ = 7 used in the simulations.

#### Interaction potentials

In our model, all potentials can be decomposed into pairwise interactions. Potentials involving shell subunits further decompose into pairwise interactions between their constituent building blocks – the excluders, attractors, ‘Top’, and ‘Bottom’ pseudoatoms. It is convenient to write the total energy of the system as the sum of 6 terms:, involving shell-shell (*U*_hh_), scaffold-shell (*U*_sh_), scaffold-cargo (*U*_sc_), scaffold-scaffold (*U*_ss_), cargo-cargo (*U*_cc_) and shell-cargo (*U*_hc_) interactions, each summed over all pairs of the appropriate type:

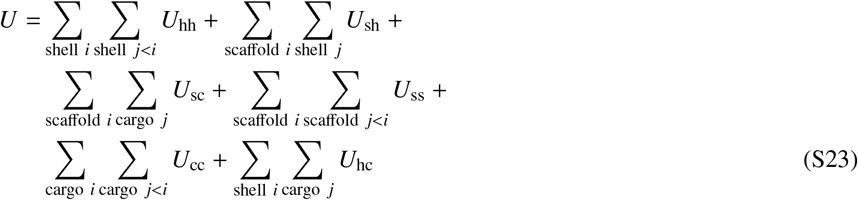

where 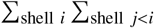 is the sum over all distinct pairs of shell subunits in the system, 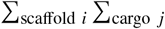 is the sum over all shell-scaffold particle pairs, etc.

##### Shell-shell interaction potentials

The shell-shell potential *U*_ss_ is the sum of the attractive interactions between complementary attractors, and geometry guiding repulsive interactions between ‘Top’ - ‘Top’, ‘Bottom’ - ‘Bottom’, and ‘Top’ - ‘Bottom’ pairs. There are no interactions between members of the same rigid body. For notational clarity, we index rigid bodies and non-rigid pseudoatoms in Roman, while the pseudoatoms comprising a particular rigid body are indexed in Greek. For subunit *i* we denote its attractor positions as {**a**_*iα*_} with the set comprising all attractors *α*, its ‘Top’ position **t**_*i*_, ‘Bottom’ position **b**_*i*_ and the ‘M’ pseudoatom at the center of the subunit in the plane of the attractors, as **m**_*i*_.

The shell-shell interaction potential between two subunits *i* and *j* is then defined as:

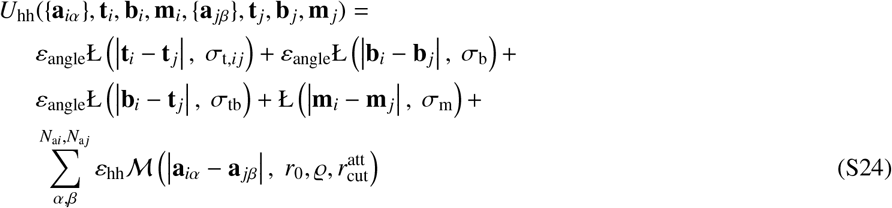

The function Ł is defined as the repulsive component of the Lennard-Jones potential shifted to zero at the interaction diameter:

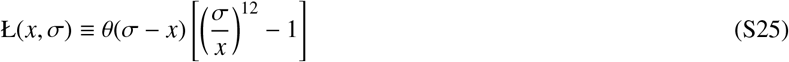

with *θ*(*x*) the Heaviside function. The function 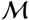 is a Morse potential:

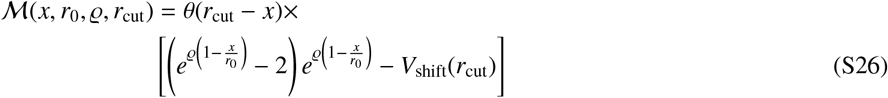

with *V*_shift_(*r*_cut_) the value of the (unshifted) potential at *r*_cut_.

The parameter *ε*_hh_ sets the strength of the shell-shell attraction at each attractor site, *N_ai_* is the number of attractor pseudoatoms in subunit *i*, and *ε*_angle_ scales the repulsive interactions that enforce the geometry.

##### Shell-shell interaction parameter values

*Attractors*: The strength of attractive interactions is parameterized by the welldepth *ε*_hh_ for a pair of attractors on hexamers as follows. Hexamer-Hexamer edge attractor pairs have a well-depth of *ε*_hh_. Because vertex attractors have multiple partners in an assembled structure, whereas edge attractors have only one, the well-depth for the vertex pairs is set to 0.5*ε*_hh_. The parameter *r*_0_ is the minimum energy attractor distance, set to 0.2, 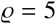 determines the width of the attractive interaction, and 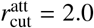 is the cutoff distance for the attractor potential.

*Repulsive interactions and simulated shell bending modulus:* The ‘Top’ and ‘Bottom’ heights, or distance out of the attractor plane, are set to *h* = 1/2. The diameter of the ‘Top’ - ‘Bottom’ interaction, which prevents subunits from binding in inverted configurations [77], is *σ*_tb_ = 1.8. A central excluder ‘M’ with effective diameter *σ*_m_ = 2.026 in the center of subunits prevents subunit overlaps. The effective diameters of the ‘Bottom’ - ‘Bottom’ interaction σ_b_ and the ‘Top’ - ‘Top’ interaction σ_t_ determine the shell spontaneous curvature radius as follows: *R*_0_ = 22, *σ*_b_ = 1.6, *σ*_t_ = 2.2; *R*_0_ = 8.0, *σ*_b_ = 1.5, *σ*_t_ = 2.3; and *R*_0_ = 4.5, *σ*_b_ = 1.4, *σ*_t_ = 2.4.

The bending modulus of *α*-carboxysomes has not been experimentally estimated, but measurements on *β*-carboxysome shells obtained *κ* ≈ 25*k*_B_*T* [142], which is thus smaller than that for a typical viral capsid. As noted above, our simulated hexamer subunits, with diameter 13 nm, are larger than the hexamer size (7 nm diameter) in a carboxysome. To maintain the characteristic elastic length scale 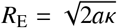 of the simulated shells approximately equal to that of *β*-carboxysome shells *R*_E_ ≈ 47 nm, despite the large size of hexamer subunits, we set *ε*_angle_ = 0.5*k*_E_*T*, which corresponds to a bending modulus of about 5-10 *k*_B_*T*[45], and thus *R*_E_ ≈ 38 – 54 nm.

##### Scaffold-shell interactions

The scaffold-shell interaction is modeled by a short-range repulsion between pairs of scaffold beads and scaffold-excluder psudoatoms on subunits representing the excluded volume, plus an attractive interaction between pairs of shell-interacting beads in scaffolds and hexamer ‘Bottom’ pseudoatoms. For subunit *i* with excluder positions {**x**_*iα*_} and ‘Bottom’ psuedoatom position **b**_*i*_, and scaffold *j* with bead positions **s**_*jβ*_ and shell-interacting beads *jγ* with positions **s**_*jγ*_, the potential is:

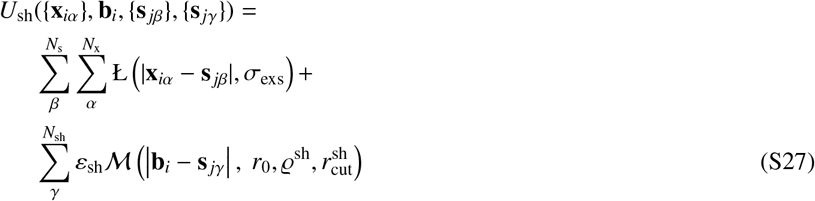

where *ε*_sh_ parameterizes the scaffold-shell interaction strength, *N*_s_ and *N*_sh_ are the total number of beads and the number of shell-interacting beads in each scaffold, *N*_x_ is the numbers of excluders on a shell subunit, *σ*_exs_ = 0.375 and *σ*_t_ = 0.375 are the effective diameters of the excluder - scaffold repulsions, 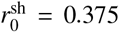 is the minimum energy attractor distance, the width parameter is 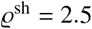, and the cutoff is set to 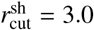.

##### Scaffold-cargo interactions

The interaction between cargo particles and scaffolds is given by

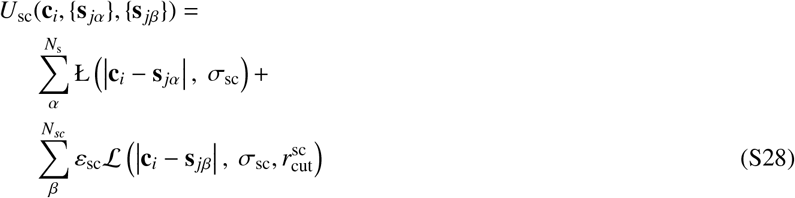

with 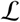 the full Lennard-Jones interaction:

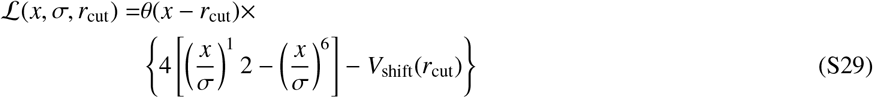

with *ε*_sc_ an adjustable parameter that sets the strength of the scaffold-cargo interaction, *N_s_* as the total number of scaffold beads, *N*_sc_ as the number of beads in the scaffold-cargo binding domain, *σ*_sc_ = 0.75 as the effective cargo-scaffold excluded volume size, and 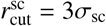 the interaction cut off length.

##### Scaffold-scaffold interactions

The scaffold-scaffold non-bonded interaction is modeled by a Lennard-Jones potential; in addition, segments occupying adjacent positions along the polymer chain interact through harmonic bonds.

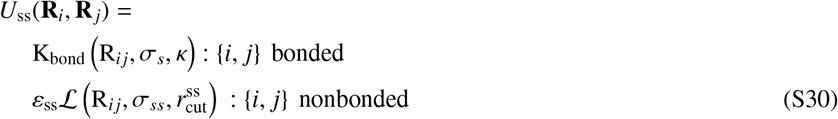

where *R_ij_* ≡ |**R**_*i*_ – **R**_*j*_| is the center-to-center distance between the scaffold beads, *ε*_ss_ is the well-depth of the Lennard-Jones (Eq. S29) potential, *σ*_ss_ = 0.5 is the effective scaffold-scaffold excluded volume size, and 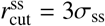 the interaction cut off length.

Bonds are represented by a harmonic potential:

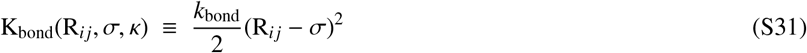

##### Cargo-cargo interactions

The cargo-cargo interaction is modeled by a short-range repulsion interaction representing the excluded volume of cargo particles, given by:

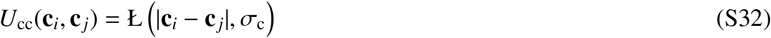

##### Shell-cargo interactions

The shell-cargo interaction is modeled by a short-range repulsion between pairs of cargo particles and cargo-excluder psudoatoms on subunits representing the excluded volume:

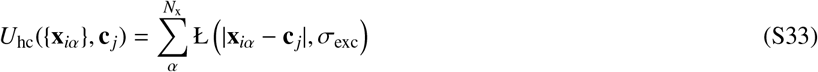

**FIG. S1.**
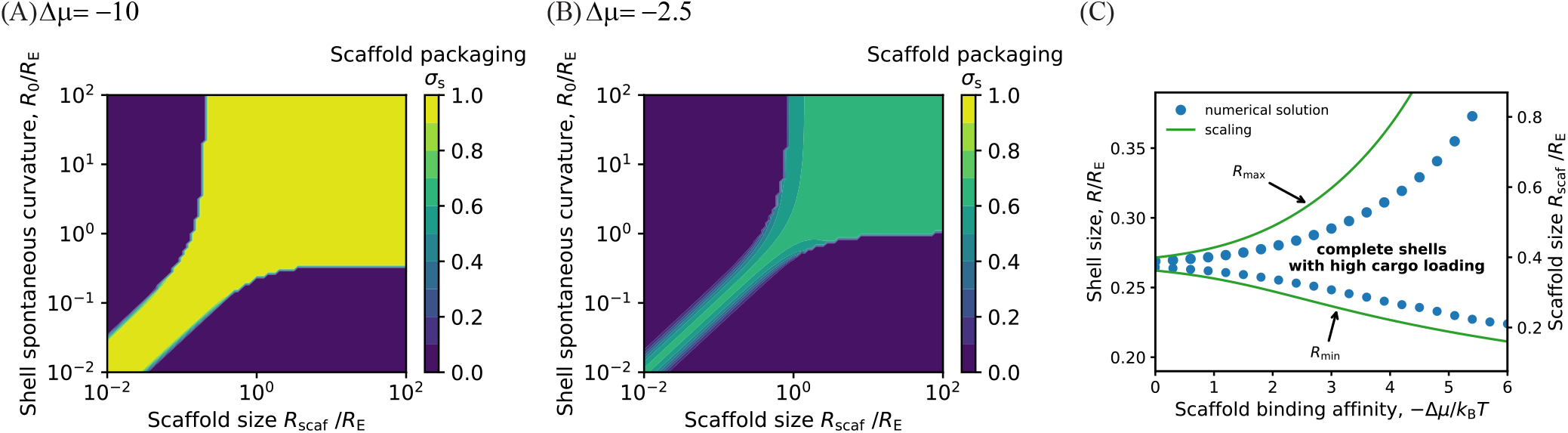
Equilibrium theory prediction for the amount of packaged scaffold *σ*_s_ as a function of *R*_0_ and *R*_scaf_. Predictions are obtained by numerically minimizing Eq. 1 with *R*_E_ = 30*r*\*, and *ν* = 0. Results are shown for **(A)** Δ*μ* = −10 and **(b)** Δ*μ* = −2.5. **(C)** Minimum and maximum shell radius (normalized by the shell elastic energy length scale *R*_E_) in the high intrinsic shell curvature regime (*R*_0_ < *R*_E_). The maximum and minimum shell radii that can form for varying scaffold length are shown, calculated by numerically minimizing Eq. 1 (symbols) and from the asymptotic analysis Eq. 11 (lines) at different scaffold binding affinities Δ*μ* for shell spontaneous curvature *R*_0_ = 8, *R*_E_ = 30 and *ν* = 0.

**FIG. S2.**
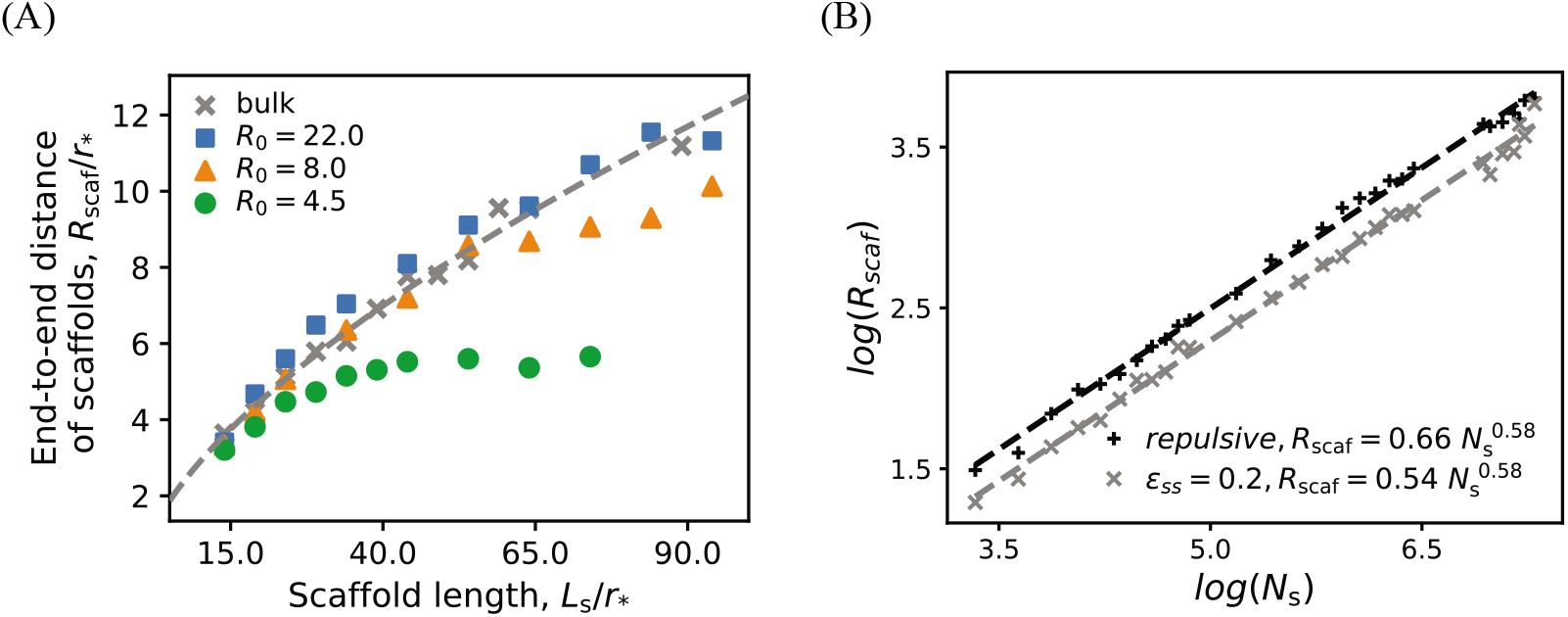
**(A)** End-to-end distance (*R*_scaf_) of scaffolds encapsulated in shells as a function of scaffold length. For comparison, the unperturbed scaffold sizes (from panel (B)) are shown as gray ‘x’ symbols and the dashed line. **(B)** End-to-end distance of isolated scaffold molecules (i.e., free in solution rather than packaged in a shell) as a function of number of segments *N*_s_, with the scaffold contour length given by *L*_s_ = *l*_k_*N*_s_ with *l*_k_ = 0.5*r*\*. The plot shows the end-to-end length measured with weak attractions between scaffold segments, *ε*_ss_ = 0.2 (as used in the assembly simulations), and for comparison, the end-to-end scaling with purely repulsive inter-segment interactions.

**FIG. S3.**
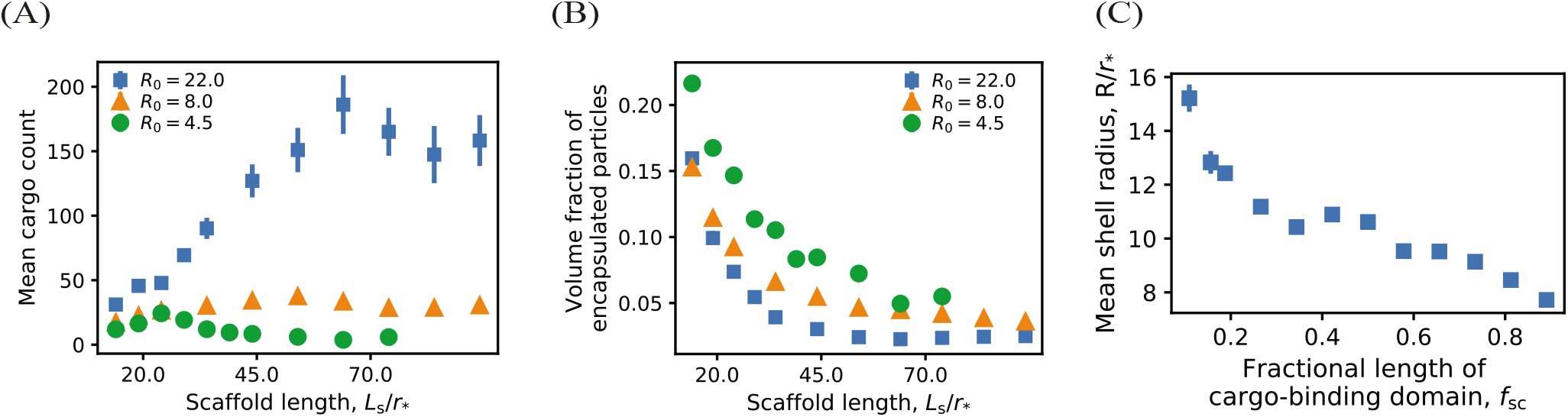
**(A)** Dependence of the number of encapsulated cargo particles on the scaffold length. **(B)** Dependence of the volume fraction of encapsulated particles (cargo and scaffolds) on the scaffold length. **(C)** Dependence of the shell size on the the scaffold cargo-binding domain fraction *f*_sc_ = *L*_sc_/*L*_s_, for *R*_0_ = 22*r*\*, *L*_s_ = 64*r*\*, and *L*_sh_ = 7*r*\*. Other parameter values for (A), (B) and (C) are as follows. Shell subunit-subunit affinities: *ε*_hh_ = 3.15 at *R*_0_ = 4.5, *ε*_hh_ = 2.85 at *R*_0_ = 8.0 and *ε*_hh_ = 2.65 at *R*_0_ = 22; scaffold-shell affinity *ε*_sh_ = 2.5, and scaffold-cargo affinity *ε*_sc_ = 1.0.

**FIG. S4.**
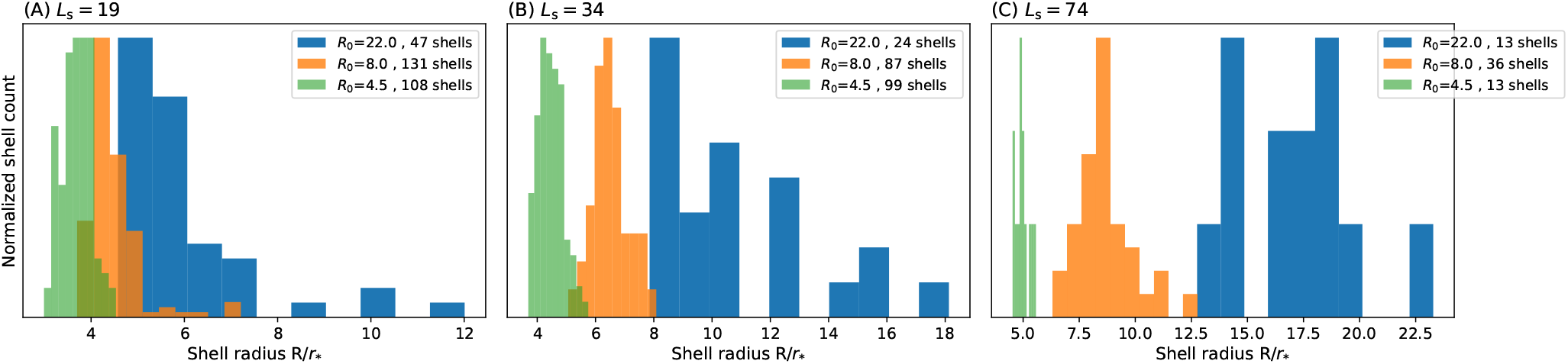
Distribution of shell sizes at three scaffold lengths and three different radii of curvature of Fig. 3.

**FIG. S5.**
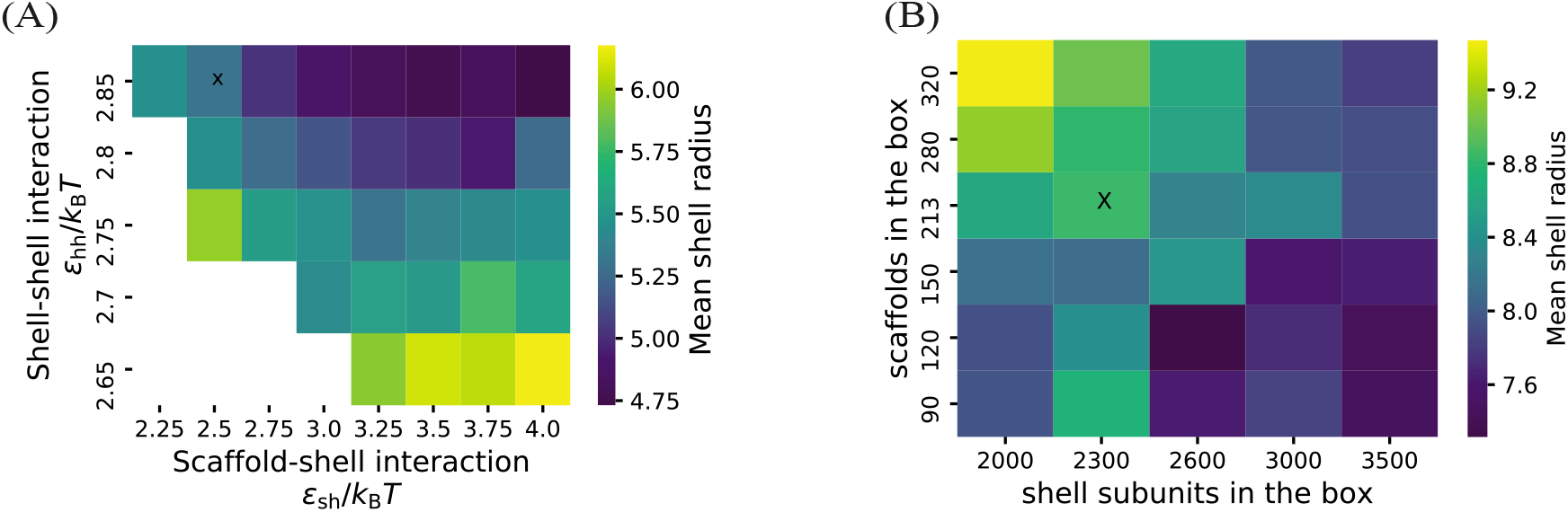
**(A)** Dependence of the mean shell radius on the scaffold-shell and shell-shell affinities. The scaffold length is *L*_s_ = 24 (*L*_sc_ = 7, *L*_sm_ = 10), the shell spontaneous curvature radius is *R*_0_ = 8.0*r*\* and the scaffold-cargo affinity is *ε*_sc_ = 1*k*_B_*T*. Note that in the large *L*_s_ limit where relatively few scaffold molecules are packaged, strong shell-shell interactions (*ε*_ss_ > 3.15 at *R*_0_ = 4.5, *ε*_ss_ > 2.85 at *R*_0_ = 8 and *ε*_ss_ > 2.65 at *R*_0_ = 22) are required for complete shell assembly. **(A)** Dependence of the mean shell radius on shell-subunit and scaffold concentrations. The scaffold length is *L*_s_ = 64 (*L*_sc_ = 7, *L*_sm_ = 50), the shell spontaneous curvature radius is *R*_0_ = 8.0*r*\*, and the scaffold-cargo affinity is *ε*_sc_ = 1.0*k*_B_*T*. In (A) and (B), the parameter values corresponding to the simulation results of Figs. (2) and (3) are marked on the plot by ‘x’ symbols.

